# Feature-based attention multiplicatively scales the fMRI-BOLD contrast-response function

**DOI:** 10.1101/2022.03.15.484428

**Authors:** Joshua J. Foster, Sam Ling

**Affiliations:** Department of Psychological and Brain Sciences, Boston University, Boston MA, 02215; Center for Systems Neuroscience, Boston University, Boston MA, 02215

**Author notes:** **Corresponding authors:** Joshua J. Foster or Sam Ling.

## Abstract

Functional MRI (fMRI) plays a key role in the study of attention. However, there remains a puzzling discrepancy between attention effects measured with fMRI and with electrophysiological methods. While electrophysiological studies find that attention increases sensory gain, amplifying stimulus-evoked neural responses by multiplicatively scaling the contrast-response function (CRF), fMRI appears to be insensitive to these multiplicative effects. Instead, fMRI studies typically find that attention produces an additive baseline shift in the blood-oxygen-level-dependent (BOLD) signal. These findings suggest that attentional effects measured with fMRI reflect top-down inputs to visual cortex, rather than the modulation of sensory gain. If true, this drastically limits what fMRI can tell us about how attention improves sensory coding. Here, we re-examined whether fMRI is sensitive to multiplicative effects of attention using a feature-based attention paradigm designed to preclude any possible additive effects. We measured BOLD activity evoked by a probe stimulus in one visual hemifield while participants (6 male, 6 female) attended to the probe orientation (attended condition), or to an orthogonal orientation (unattended condition), in the other hemifield. To measure CRFs in visual areas V1-V3, we parametrically varied the contrast of the probe stimulus. In all three areas, feature-based attention increased contrast gain, improving sensitivity by shifting CRFs towards lower contrasts. For a subset of visual eccentricities, we also found an increase in response gain, an increase in the responsivity of the CRF. These results provide clear evidence that the fMRI-BOLD signal is sensitive to multiplicative effects of attention.

**Significance Statement:** Functional MRI (fMRI) plays a central role in the study of attention because it allows researchers to precisely and non-invasively characterize the effects of attention throughout the brain. Electrophysiological studies have shown that attention increases sensory gain, amplifying stimulus-evoked neural responses. However, a growing body of work suggests that the BOLD signal that is measured with fMRI is not sensitive to these multiplicative effects of attention, calling into question what we can learn from fMRI about how attention improves sensory codes. Here, using a feature-based attention paradigm, we provide evidence that the BOLD signal can pick up multiplicative effects of attention.

## Introduction

Attention plays a central role in perception, prioritizing processing of relevant information. One way that attention improves neural coding is by increasing *sensory gain*, selectively amplifying stimulus-evoked responses in sensory areas (Hillyard et al., 1998; Carrasco, 2011). For instance, attention increases the spike rate of individual neurons in non-human primates (McAdams and Maunsell, 1999; Treue and Martinez-Trujillo, 1999) and the amplitude of visually evoked potentials measured with EEG in humans (van Voorhis and Hillyard, 1977; Morgan et al., 1996). More recent work has characterized how the effect of attention scales with stimulus drive by varying stimulus contrast to measure contrast-response functions (CRFs). Unit-recording and EEG studies that have taken this approach have found that attention *multiplicatively* scales CRFs, either vertically scaling the CRF, increasing responsivity (*response gain*, Fig. 1a), or shifting the CRF towards lower contrasts, increasing sensitivity (*contrast gain*, Fig. 1b) (Reynolds et al., 2000; Martínez-Trujillo and Treue, 2002; Williford and Maunsell, 2006; Kim et al., 2007; Itthipuripat et al., 2014a, 2014b, 2019; Foster et al., 2021). These sensory gain effects have served as the empirical backbone for computational models of attention (Reynolds and Heeger, 2009).

**Figure 1.**
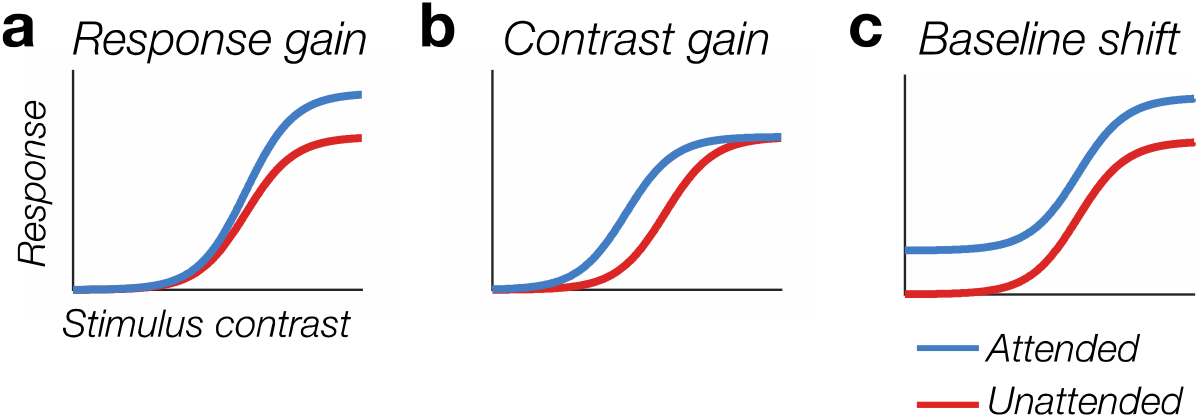
Attentional modulations of contrast-response functions (CRFs). Attention can modulate CRFs in three ways. (a) Response gain: attention increases the amplitude of the CRF, increasing responsivity. (b) Contrast gain: attention shifts the CRF, increasing sensitivity. (c) Baseline shift: attention produces an additive increase in the CRF.

Functional MRI (fMRI) has long played a central role in attention research, allowing researchers to precisely and non-invasively measure the effects of attention throughout the human brain. Like electrophysiological studies, fMRI studies report larger responses to attended stimuli than to unattended stimuli (Carrasco, 2011). It is often assumed that these effects reflect an amplification of stimulus-driven activity. However, fMRI studies of attention that have measured CRFs reveal a qualitatively different pattern than electrophysiological studies. Unlike the multiplicative effects seen in electrophysiology, fMRI studies have found that attention produces an *additive* baseline shift in the BOLD signal (Fig. 1c) (Somers and McMains, 2005; Buracas and Boynton, 2007; Murray, 2008; Pestilli et al., 2011; Sprague et al., 2018; Itthipuripat et al., 2019; but see Li et al., 2008). Because a baseline shift is independent of stimulus strength, and is seen in the absence of visual input, this pattern suggest that attentional effects measured with fMRI reflect top-down input to visual cortex, rather than a multiplicative modulation of stimulus-driven activity (Murray, 2008; Itthipuripat et al., 2014a). Understanding whether the BOLD signal is sensitive to multiplicative attention effects is critical for understanding the degree to which fMRI can be used to study attention: if the BOLD signal is not sensitive to modulations of stimulus-driven activity, then this drastically limits what fMRI can tell us about how attention improves sensory codes.

Although studies of spatial attention suggest that fMRI is insensitive to multiplicative effects of attention, a seminal study of *feature-based attention* suggests otherwise. Saenz and colleagues (2002) measured the effect of feature-based attention on the BOLD response to a spatially unattended probe stimulus. They found that attention to a specific feature (e.g. red or upward motion) increased the BOLD response to the probe stimulus when it matched the attended feature. This effect appears to reflect a multiplicative amplification of the probe-evoked response rather than a baseline shift. Although feature-based attention does produce baseline shifts in neural activity (Chelazzi et al., 1993; Serences and Boynton, 2007; Stokes et al., 2009), these baseline shifts are *feature specific*, selectively increasing the baseline activity of neurons tuned for the attended feature. Thus, the baseline activity of the entire neuronal population, when aggregating across populations of neurons with different feature preferences, is not expected to vary with the attended feature. Although Saenz et al.’s (2002) results hint at a multiplicative effect of feature-based attention, no study has assessed the effect of feature-based attention on the full CRF, which is essential to distinguish between different kinds of attentional modulation because all mechanisms can account for an increase in the response at a single contrast level (Fig. 1).

Here, we provide that definitive test. We manipulated feature-based attention in one hemifield, and measured responses to a probe stimulus in the other hemifield, exploiting the well-established spatial spread of feature-based attention (Treue and Martinez-Trujillo, 1999; Saenz et al., 2002). Critically, however, we parametrically manipulated the contrast of the probe stimulus to measure the CRF in visual cortex, using a recently developed paradigm that captures compressive nonlinearities in the BOLD CRF (Vinke et al., 2022). In doing so, we found robust multiplicative effects of feature-based attention on the CRF throughout early visual cortex. Throughout early visual cortex, feature-based attention increased *contrast gain*, shifting CRFs leftwards. For V3 and a subset of eccentricities in V2, we also found an increase in response gain. These results provide clear, positive evidence that BOLD responses are sensitive to multiplicative effects of feature-based attention on stimulus-driven responses.

## Materials and Methods

### Participants

Thirteen healthy volunteers (7 male and 6 female, mean age = 27.8 years, SD = 6.3) participated in the study. Participants were between 19 and 42 years old, and reported normal or corrected-to-normal visual acuity. Our target sample size was 12 participants. We replaced one participant because of very weak activation of visual cortex in the independent visual localizer scan (see Visual Localizer). Thus, we were not confident that the signal-to-noise ratio would be adequate to measure CRFs for this subject. The decision to exclude this subject was made solely on the basis of their visual localizer data, and we did not analyze their data in the feature-based attention task. Thus, our final sample comprised 12 participants (6 male and 6 female, mean age = 27.6 years, SD = 6.6). All participants provided written informed consent, and completed a screening form to ensure that they had no MRI-related contraindications. Participants received monetary compensation for their time ($15/hr for behavioral sessions and $40/hr for MRI sessions), except for two participants who were the authors of the study. All procedures were approved by the Boston University Institutional Review Board.

### Apparatus and stimuli

Stimuli were generated using Matlab (MathWorks) and the Psychophysics Toolbox (Brainard, 1997; Pelli, 1997). In the MRI scanner, participants viewed stimuli on a gamma-corrected, rear-projection screen (ProPixx DLP LED, VPixx Technologies; refresh rate: 60 Hz; resolution: 1024 × 768 pixels; viewing distance: ∼99 cm) that was mounted inside the scanner bore. In behavioral sessions, participants were seated at a desk with their chin on a padded chin rest and viewed stimuli on a gamma-corrected LCD monitor (Display++, Cambridge Research Systems Limited; refresh rate: 100 Hz; 1440 × 1080 pixels; viewing distance: ∼135 cm). In all sessions, the display was the only light source in the testing room. Participants responded using a two-button response box in the scanner, and using the “1” and “2” keys on the numberpad of a standard keyboard in behavioral sessions.

Stimuli were presented on a mean-luminance background (∼150 cd/m^2^). A light-gray fixation dot (0.2°) was presented in the center of the display. The stimuli were three wedge-shaped gratings (2.2 cycles/°, inner edge: 1.0°, outer edge: 8.5°, SD of Gaussian roll-off: 0.1°; see Fig 2a). The wedges in each hemifield spanned 70° of polar angle above and below the horizontal meridian. The display was divided into a *relevant hemifield* that contained task-relevant stimuli, and a *probe hemifield* that contained an irrelevant probe stimulus to evoke visual responses (counterbalanced across observers; Fig. 2a). In the relevant hemifield, the wedge was divided into two gratings, one in the upper visual field and one in the lower visual field, separated by a 0.5° gap. On each trial of the task, one of these gratings was oriented horizontally, and the other oriented vertically, with each orientation appearing in the upper and lower position equally often. In the probe hemifield, a horizontal probe grating spanned both the upper and lower visual fields. The spatial phase of each grating randomly updated at 10 Hz, with the two horizontal gratings moving in phase together. The two gratings in the relevant hemifield were always presented at 16% Michelson contrast. The probe stimulus was presented at 10 different contrast levels (0, 2.67, 4.0, 5.33, 8.0, 16, 32, 48, 64, 96%), which allowed us to measure probe-evoked CRFs.

**Figure 2.**
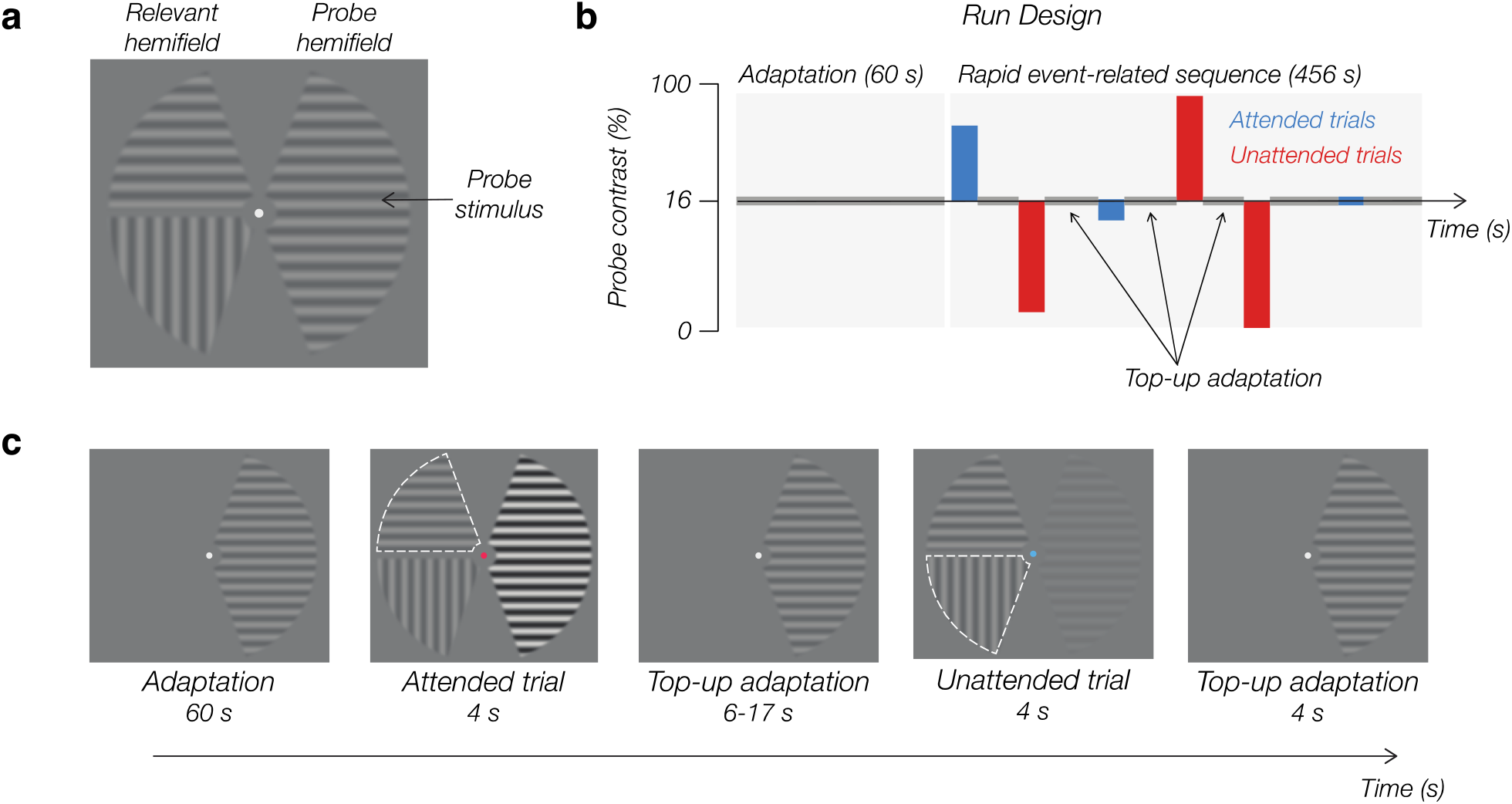
Feature-based attention task. (a) Stimulus configuration. On each trial of the task, observers were cued to attend to one of two gratings in the relevant hemifield, while we measured responses evoked by a probe stimulus in the other hemifield. (b) Each fMRI run began with a 60-s adaptation period during which the probe grating was presented at 16% contrast to adapt visual neurons to this contrast level. Following the initial adaptation period, 4-s trials were interleaved with periods of top-up adaptation. The contrast of the probe stimulus was parametrically varied across trials (0-96%). (c) Illustration of the task sequence. On each trial, the color of the fixation dot cued observers to attend one of the two gratings in the relevant hemifield (marked by white dotted outline that was not visible to observers). We manipulated feature-based attention by varying whether observers attended the horizontal or vertical grating in the relevant hemifield. On attended trials, observers attended to the horizontal grating, which matched the orientation of the probe. On unattended trials, observers attended the vertical grating, which did not match the orientation of the probe. Note: The spatial frequency of the gratings in this figure has been reduced for clarity.

### Task procedure

Participants completed a rapid event-related task (Fig. 2b and 2c). We used a contrast-adaptation procedure to promote compressive nonlinearities in CRFs (Vinke et al., 2022, see Results, Task and Behavior). Each run began with a 60-s adaptation period, during which the probe stimulus was presented at 16% contrast (the adapting contrast). The initial adaptation period was followed by a 456-s rapid event-related trial sequence, with 4-s trials interleaved with top-up adaptation periods (6-17 s) to maintain the adaptation state (Fig. 2b). The timing of the trials and order of conditions (20 total: 2 attention conditions × 10 probe contrasts) was sequenced using the Optseq2 schedule optimization tool (Dale, 1999). Participants completed 9-10 runs of the task in the scanner. Each run included 40 trials (2 trials per condition). Therefore, we obtained 18-20 trials per condition for each participant.

During each trial, two gratings were presented in the relevant hemifield, and the contrast of the probe stimulus changed (Fig. 2c). The color of the fixation dot (pink or blue) cued the observer to attend the upper or lower grating in the relevant hemifield (with color-location mapping counterbalanced across observers). The color cue onset 0.5 s before the 4-s trial period and remained on-screen until the end of the trial. On half of trials, the cued grating was horizontal, such that the attended orientation matched the orientation of the probe stimulus (*attended* trials). On the other half of trials, the cued grating was vertical, such that the attended orientation did not match the probe stimulus (*unattended* trials). The observer’s task was to monitor the cued grating for targets, which were occasional, brief (100 ms) reductions in the grating’s spatial-frequency. On each trial, zero, one, or two target reductions could occur. Targets never appeared during the first 0.7 s or the final 0.2 s of the trial. On trials with two targets, the targets were always separated by at least 0.6 s to avoid poor detection of the second target due to the attentional blink (Raymond et al., 1992; Duncan et al., 1994), and the second target always occurred during the final second of the trial to ensure that observers needed to remain engaged with the task for the full trial duration. Similar spatial frequency perturbations occurred in the two unattended gratings to encourage observers to selectively attend to the cued grating. For the probe grating, these perturbations were an increase in spatial frequency rather than a decrease, which ensured that on attended trials, observers could not perform the task by simply monitoring for a relative decrease in spatial frequency between the cued horizontal grating and the collinear probe grating in the other hemifield. Following the trial period, observers reported how many targets they detected with a button press (no response = zero targets; left button = one target; right button = two targets). Responses needed to be made within 2 s of the end of the trial. At the end of the response window, the fixation dot changed turned green or red for 0.5 s to indicate whether the response was correct or incorrect, respectively.

### Staircase procedure

Prior to the MRI session, participants completed 1-2 behavioral practice sessions (∼1.5 hours each) to learn the task and to perform initial staircasing to equate difficulty between the attended and unattended conditions. For each condition, we ran separate staircases for trials in which the attended wedge was in the upper or lower visual field to account for potential differences in performance between locations. We titrated difficulty by adjusting the size of the spatial frequency decrement with a weighted up/down procedure: after a correct response, we decreased the size of the spatial frequency decrement by 5%; after an incorrect response, we increased the size of the spatial frequency decrement by 17.6%. The staircases operated continuously throughout the behavioral session(s) and the MRI session. During the behavioral sessions, participants completed staircasing runs of the task until performance stabilized in each staircase (7-15 runs, M = 11.6, SD = 2.0). This procedure successfully equated accuracy across conditions during the MRI session: mean accuracy was 76.6% (SD = 2.4) in the attended condition and 77.0% (SD = 2.0) in the unattended condition.

### Eye tracking

We monitored eye position in the scanner using an MR-compatible EyeLink 1000 Plus infrared eye tracker (SR Research). Eye position was sampled at 500 Hz, and we calibrated the eye tracker as needed at the start of each run of the task. After removing samples contaminated by blinks, we calculated the mean eye position (relative to the fixation dot) during the 4-s trial period for each condition. The eye tracking data confirmed that observers did a good job of maintaining fixation. We observed a very small bias (< 0.1°) in vertical eye position toward the location of the cued grating: mean vertical eye position was 0.069° (SEM = 0.097) when the upper grating was cued and –0.046° (SEM = 0.116) when the lower grating was cued, *t*(11) = 3.29, *p* = 0.007, but horizontal eye position did not vary a function of location of the cued location (upper cued: M = 0.157°, SEM = 0.055, lower cued: M = 0.166°, SEM = 0.048, *t*(11) = –0.71, *p* = 0.491). More importantly, however, eye position did not differ between the attended and unattended conditions: mean vertical eye position (attended: M = 0.014°, SEM = 0.106; unattended: M = 0.010°, SEM = 0.106, *t*(11) = 0.49, *p* = 0.631) and horizontal eye position (attended: M = 0.160°, SE = 0.052; unattended: M = 0.163°, SEM = 0.050, *t*(11) = -0.28, *p* = 0.785) were near-identical across conditions. Thus, any effects of feature-based attention cannot be attributed to differences in eye position.

### Visual localizer

Participants performed one run of a visual localizer task to identify voxels that were responsive to the wedge stimuli used in the feature-based attention task. The wedge stimuli presented in the localizer task were identical to those in the feature-based attention task (see Apparatus and stimuli), except that all three gratings were horizontally oriented and were presented at full contrast. We alternated 16-s periods of blank and stimulation eight times, followed by a final 16-s blank period (272 s total). To encourage fixation and maintain alertness levels, participants monitored the fixation dot for 300-ms decrements in luminance that occurred every 3-5 seconds, reporting these target events with a button-press. The hit rate was above 90% for all participants (mean = 98.1%, SD = 2.4). We analyzed the visual localizer data using a standard general linear model (GLM) analysis in Freesurfer (Fischl, 2012).

### MRI data acquisition

All neuroimaging data was acquired at the Boston University Cognitive Neuroimaging Center, on a 3T Siemens Prisma scanner (Siemens Healthcare, Erlangen, Germany) using the vendor’s 64-channel headcoil. A whole-brain anatomical scan was acquired using a T1-weighted magnetization-prepared rapid gradient multi-echo MPRAGE sequence (van der Kouwe et al., 2008), with the following parameters: 1.0 mm^3^ voxels, 176 sagittal slices, TR = 2530 ms, TEs = 1.67, 3.55, 5.41, and 7.27 ms, TI = 1100 ms, flip angle = 7°, FOV = 256 mm, GRAPPA (Griswold et al., 2002) acceleration = 2. BOLD data were collected via a T2*-weighted echo-planar imaging (EPI) pulse sequence that employed multiband RF pulses and Simultaneous Multi-Slice (SMS) acquisition (Feinberg et al., 2010; Moeller et al., 2010; Setsompop et al., 2012; Xu et al., 2013; Cauley et al., 2014). For the task runs, the scan parameters were: 2.0 mm^3^ voxels, 70 interleaved axial-oblique slices (25 degrees toward coronal from ACPC alignment), TR = 1000 ms, TE = 30 ms, FA = 64°, FOV = 208 mm, SMS factor = 5, GRAPPA acceleration = 2. The SMS-EPI acquisition used the CMRR-MB pulse sequence from the University of Minnesota.

### Anatomical analysis

We analyzed whole-brain T1-weighted anatomical data using the standard ‘recon-all’ pipeline provided by FreeSurfer (Fischl, 2012) to generate cortical-surface reconstructions, which enabled surface-based registration of functional data to structural data. This allowed us to align population receptive field (pRF) data (see *Population receptive fields*) to the functional volume space for the feature-based attention task.

### fMRI preprocessing

Functional BOLD time-series were corrected for echo-planar imaging (EPI) distortions using a reverse phase-encoded method (Andersson et al., 2003) implemented in FSL (Smith et al., 2004). The fieldmap-corrected data were preprocessed with FS-FAST (Fischl, 2012) to apply standard motion-correction procedures, Siemens slice timing correction, and boundary-based registration between functional and anatomical spaces (Greve and Fischl, 2009). No spatial smoothing was applied. To ensure good voxel-wise correspondence of functional data across runs, we applied robust rigid registration (Reuter et al., 2010) using the middle time point of each run. Additionally, for runs of the feature-based attention task, we excluded the first 60 seconds of data that corresponded to the initial adaptation period (Fig 2b and 2c; see Task procedure), before the time-series for each voxel was linearly detrended, high-pass filtered (low cutoff = 0.01 Hz), and converted to z-scores. We concatenated the data for all runs of the feature-based attention task. For one participant, we excluded one run of the feature-based attention task due to excessive head motion.

### Population receptive fields

pRF mapping was conducted for each participant in a separate scan session. Observers completed 3-5 scans of each of two types of mapping runs: (1) expanding/contracting ring and bar sweep stimuli; and (2) rotating wedge stimuli. The stimuli were composed of a pink noise background with colored objects and faces of varying spatial scale (refreshed at 15 Hz) on mean luminance background (Kay et al., 2013), as used in the Human Connectome Project 7T Retinotopy dataset (Benson et al., 2018). The data were analyzed with the analyzePRF toolbox for Matlab, which implements the compressive spatial summation pRF model (Kay et al., 2013). Only voxels within the cortical ribbon of the occipital lobe were included in pRF analysis, which were identified using a visual area network label generated using an intrinsic functional connectivity atlas (Yeo et al., 2011). The results of the pRF analysis were used to manually draw region-of-interest (ROI) labels for early visual areas V1, V2 and V3.

### Voxel selection

We used the estimated pRFs in conjunction with the visual localizer (see Visual localizer) data to select voxels for analysis in the feature-based attention task. For each ROI, we excluded voxels whose pRF was centered outside the apertures of the wedge stimuli used in the FBA task, whose pRF model fit was poor (R^2^ < 10%), or whose size was unreasonably small (RF < 0.1°). Of the remaining voxels, we identified the 50% most visually responsive voxels based on the localizer data (see Visual localizer) for analysis. On average, this procedure left 160 ± 42 (SD) voxels in V1, 147 ± 25 in V2, and 129 ± 11 voxels in V3 for each hemisphere.

### fMRI data analysis

We used a general linear model (GLM) to estimate the deconvolved hemodynamic response function (HRF) in each condition. Specifically, we modeled time points from 0 to 24 s (25 time points) relative to trial onset using a finite impulse response approach, and included run-specific terms to account for any baseline differences between runs. Given each voxel’s time-series, we solved a GLM using standard linear regression to obtain 25 weights for each condition, providing an estimate of the voxel’s HRF in each condition.

### Measuring contrast-response functions

We measured CRFs by taking the average of the deconvolved HRF over a fixed window 3–9 seconds after stimulus onset for each probe contrast level, per attention condition. We fit the CRFs in each condition with a Naka-Rushton of the form:

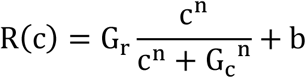

Where *R* is the measured BOLD response, and *c* is stimulus contrast. The function had four free parameters: baseline (*b*), which determines the offset of the function from zero, response gain (*G*_*r*_), which determines the amplitude of the function above baseline, contrast gain (*G*_*c*_), which determines the semi-saturation point of the function, and an exponent *(n)* that determines the slope of the function. We used the “scipy.optimize.curve_fit” function from SciPy (Virtanen et al., 2020) to fit the Naka-Rushton function to measured CRFs. We restricted the *b* parameter to be between -10 and 10, *G*_*r*_ to be between 0 and 10 (with 10 being a value that far exceeds the observed amplitudes of the CRFs), *G*_*c*_ to be between 0 and 100% contrast, and *n* to be between 0.1 and 10.

As Itthipuripat et al. (2019) have pointed out, in the absence of a fully saturating CRF, it is possible to obtain unrealistically large estimates of *G*_*r*_ when the best-fitting function saturates outside the range of possible contrasts (i.e. 0–100%). If the best fit function saturates above 100% contrast, the maximum value of the function can well exceed the largest measured response. Thus, following Itthipuripat et al. (2019), rather than reporting *G*_*r*_ and *G*_*c*_, we instead obtained a measure of response gain, *R*_*max*_, by calculating the amplitude of the best-fit Naka-Rushton function at 100% contrast and subtracting the baseline (i.e., *R*_*max*_ = *R*(100) – *b*), and a measure of contrast gain, *C*_*50*_, by calculating the contrast at which the function is halfway between baseline and the response at 100% contrast.

We used a subject-level resampling procedure to test for differences in the parameters of the fitted Naka-Rushton function across conditions. We drew 10,000 bootstrap samples, each containing N - 1 many subjects sampled with replacement, where N is the sample size. We drew samples of size N -1 to correct for bias toward narrow confidence intervals in small samples (Hesterberg, 2011). For each bootstrap sample, we fitted Naka-Rushton function to the amplitude of mean CRFs across subjects in the bootstrap sample. We calculated the difference for the parameter between the attended and unattended conditions for each bootstrap sample, which yielded a distribution of 10,000 values. We tested whether these difference distributions significantly differed from zero in either direction by calculating the proportion of values > or < 0, and doubling the smaller value to obtain a 2-sided *p* value.

### Data and code availability

All data and code will be made available on Open Science Framework at the time of publication.

## Results

### Task and behavior

Observers performed a feature-based attention task (Fig. 2; see Materials and Methods, Task procedures). We manipulated feature-based attention in one visual hemifield (the *relevant hemifield*), and measured fMRI-BOLD responses to a spatially unattended probe stimulus in the other hemifield (the *probe hemifield*). On each trial, observers viewed three flickering gratings (with spatial phase randomly updating at 10 Hz): a pair of gratings (one vertical, one horizontal) in the relevant hemifield, and a horizontal probe grating in the opposite hemifield (Fig 2a). The color of the fixation dot cued observers to covertly attend to one of the two gratings in the relevant hemifield (while maintaining fixation, see Materials and Methods, Eye tracking). On half of the trials, observers were cued to attend the horizontal grating in the relevant hemifield that matched the orientation of the probe grating (*attended trials*). On the other half of trials, observers were cued to attend the vertical grating (*unattended trials*). Thus, we could measure responses evoked by the irrelevant probe stimulus, while feature-based attention was manipulated in the relevant hemifield. During each trial of the task, observers monitored the cued grating for decrements in spatial frequency, reporting whether they detected zero, one, or two decrements. Observers practiced this task outside the scanner for several hours prior to the fMRI session. We used a staircasing procedure to adjust task difficulty (Materials and Methods, Staircase procedure), which ensured that accuracy was equated between the attended (M = 76.6%, SD = 2.4) and unattended (77.0%, SD = 2.0) conditions during the fMRI session.

To measure the full CRF in each attentional condition, we parametrically varied the contrast of the probe stimulus (0–96%). One challenge for measuring CRFs with fMRI is that the BOLD response often appears to scale linearly with stimulus contrast (e.g., Boynton et al., 1999; Murray, 2008). This is rather unlike CRFs that are typical of individual neurons, which are characterized by a compressive nonlinearity, saturating at higher contrasts (Albrecht and Hamilton, 1982; Williford and Maunsell, 2006). To overcome this challenge, we took advantage of a contrast-adaptation procedure, which recent work from our lab has shown yields BOLD CRFs that show a compressive nonlinear response, more closely mirroring the known properties of CRFs measured at the level of individual neurons (Vinke et al., 2022). Vinke and colleagues (2022) proposed that linear BOLD CRFs measured without contrast adaptation may reflect the fact that the fMRI measurements aggregate across populations of neurons that vary widely in their semi-saturation points (Fig. 3a). Unit recording studies have provided clear evidence that contrast adaptation modulates contrast gain, shifting the semi-saturation point of neurons toward the adapting contrast (Ohzawa et al., 1982; Sclar et al., 1989). Thus, contrast adaptation may yield non-linear CRFs measured with fMRI by bringing the CRFs of the neuronal population into closer alignment (Fig. 3b). Indeed, Vinke et al. (2022) found that when observers were adapted to 16% contrast stimulus, BOLD CRFs in early visual cortex showed a compressive nonlinearity, saturating at higher contrasts. Following Vinke and colleagues (2022), we began each run of the task with a 60-s adaptation period, during which observers were adapted to the probe stimulus at 16% contrast (the adapting contrast), and the trials were interleaved with top-up adaptation periods to maintain the adaptation state (Fig 2b and 2c; c.f. Gardner et al., 2005; Vinke et al., 2022).

**Figure 3.**
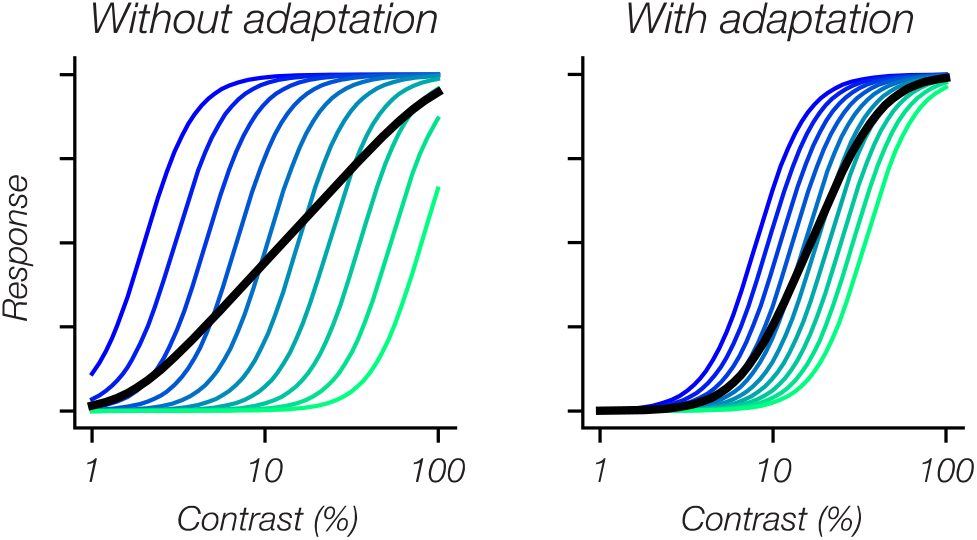
Illustration of how contrast adaptation may promote non-linear CRFs measured with fMRI. (a) fMRI measurements aggregate across many neurons that may vary considerably in their semi-saturation point without adaptation (colored curves), which yield a near-linear average CRF (black curve). (b) Unit recordings have shown that contrast adaptation shifts the semi-saturation point CRFs of neurons toward the adapting contrast. Therefore, adaptation may bring the CRFs of individual neurons into closer alignment, yielding a non-linear CRF when aggregating across the population.

### Spatial attention within the relevant hemifield modulates cortical responses

Spatial attention increases baseline BOLD activity in visual cortex (e.g. Tootell et al., 1998; Buracas and Boynton, 2007; Murray, 2008). Therefore, before examining the effects of feature-based attention, we first sought to verify that observers spatially attended to the cued grating in the relevant hemifield. To this end, we examined whether the BOLD response in the hemisphere contralateral to the relevant hemifield varied depending on which grating (upper or lower) was cued. Of the visually responsive voxels (Materials and Methods, Voxel selection), we identified voxels whose pRF center fell within the upper or lower grating in the relevant hemifield, so that we could measure voxel responses when the grating they respond to was cued or uncued (Fig. 4). We found that voxel responses were larger when their preferred grating was cued than when it was uncued (two-sided paired t-tests, V1: *t*(11) = 17.15, *p <* 0.001; V2: *t*(11) = 12.83, *p <* 0.001, V3: *t*(11) = 11.84, *p <* 0.001), confirming that observers deployed spatial attention to the cued grating.

**Figure 4.**
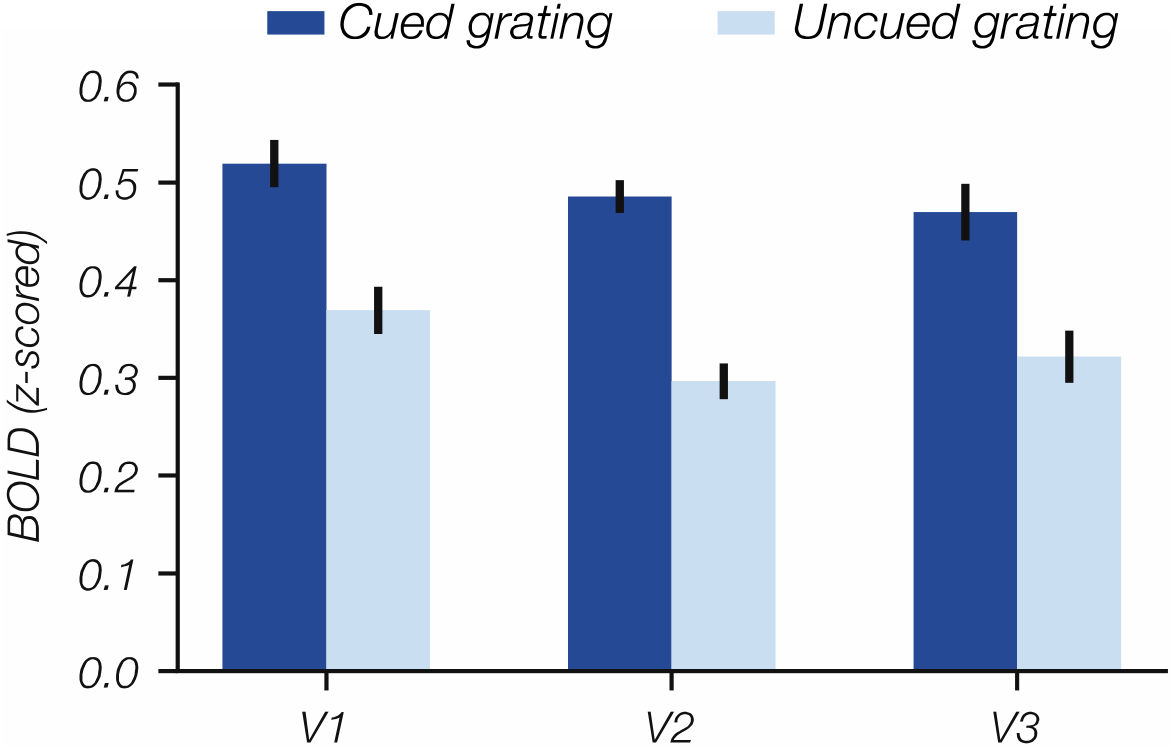
Spatial attention modulated cortical responses to stimuli in the relevant hemifield. Using pRF estimates, we identified voxels in V1-V3 that responded to each of the gratings (upper and lower) in the relevant hemifield. The bar plot shows the response measured 3-9 s after trial onset for voxels responsive to the cued grating (dark bars) and for voxels responsive to the uncued grating (light bars) for visual areas V1, V2, and V3. In all three areas, we observed greater activation of voxels selective for the cued grating than of voxels selective for the uncued grating. Error bars show ± 1 SEM across subjects.

### Feature-based attention multiplicatively scales probe-evoked responses

Having verified that observers attended the cued grating, we turned to our main question: does feature-based attention multiplicatively modulate the BOLD CRF evoked by the probe stimulus? To measure CRFs in visual areas V1-V3, we deconvolved HRFs for stimulus-responsive voxels contralateral to the probe grating as a function of probe contrast and attention condition (Fig. 5a). Figure 5b shows CRFs for the attended and unattended conditions for each of the three visual areas. To characterize the effect of feature-based attention, we fit a Naka-Rushton function to the CRFs measured in each condition (Fig 5b; Materials and Methods, Measuring contrast-response functions). For each attention condition, we estimated four parameters of the Naka-Rushton function: a baseline parameter (*b*), which determines the additive offset of the function from zero; a response gain parameter (*R*_*max*_), which determines how much the function rises above baseline, a contrast gain parameter (*C*_*50*_); which measures the semi-saturation point of the function, with changes in *C*_*50*_ producing horizontal shifts of the CRF; and a slope parameter (*n*), which determines how steeply the slope rises (c.f. Itthipuripat et al., 2019; Foster et al., 2021). Table 1 shows the Naka-Rushton parameter estimates for the attended and unattended conditions. We found no evidence for an additive baseline shift (a change in *b*). Instead, in all three visual areas, feature based attention increased contrast gain, shifting the CRFs to the left (a decrease in *C*_*50*_). In V3, we also found that feature-based attention increased response gain (an increase in *R*_max_). These results provide clear positive evidence that the BOLD signal measured with fMRI is sensitive to multiplicative changes in sensory gain due to feature-based attention.

**Table 1.** Naka-Rushton parameter estimates and resampling statistics results corresponding to the whole-ROI analysis shown in Figure 5c.

| ROI | Attended (mean ± bootstrapped SEM) | | | |
| --- | --- | --- | --- | --- |
| | Unattended (mean ± bootstrapped SEM) | | | |
|  | *p* value |  |  |  |
| | $R_{\text{Max}}$ | $C_{50}$ | $b$ | $n$ |
| V1 | $0.72 \pm 0.05$ | $28.19 \pm 3.04$ | $-0.24 \pm 0.05$ | $0.87 \pm 0.08$ |
| | $0.68 \pm 0.05$ | $39.27 \pm 4.12$ | $-0.24 \pm 0.04$ | $1.18 \pm 0.13$ |
| | $p = 0.240$ | $p < 0.001^{***}$ | $p = 0.986$ | $p < 0.001^{***}$ |
| V2 | $0.39 \pm 0.04$ | $21.53 \pm 3.70$ | $-0.16 \pm 0.03$ | $1.32 \pm 0.29$ |
| | $0.35 \pm 0.04$ | $36.29 \pm 5.44$ | $-0.18 \pm 0.02$ | $1.42 \pm 0.27$ |
| | $p = 0.112$ | $p < 0.001^{***}$ | $p = 0.465$ | $p = 0.730$ |
| V3 | $0.35 \pm 0.03$ | $22.80 \pm 3.38$ | $-0.11 \pm 0.03$ | $1.54 \pm 0.32$ |
| | $0.30 \pm 0.03$ | $35.01 \pm 4.73$ | $-0.12 \pm 0.02$ | $1.71 \pm 0.31$ |
| | $p = 0.025^*$ | $p < 0.001^{***}$ | $p = 0.477$ | $p = 0.566$ |

\* denotes $p$ -values $< 0.05$ and \*\*\* denotes $p$ -values $< 0.001$

**Figure 5.**
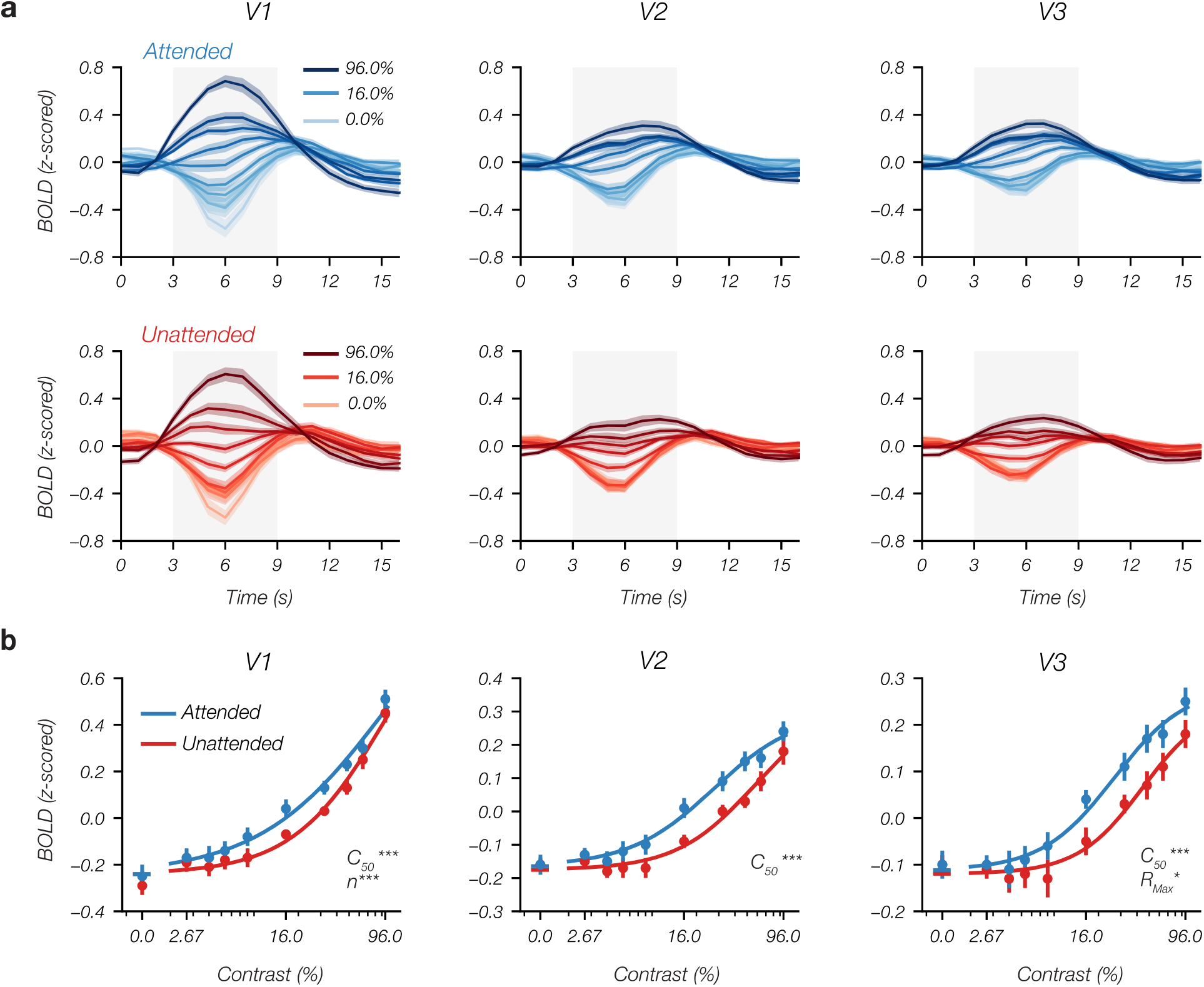
The effect of feature-based attention on probe-evoked CRFs. (a) Deconvolved HRF from stimulus-activated voxels contralateral to the probe stimulus in V1-V3 as a function of probe contrast in the attended (upper) and unattended (lower) conditions. To obtain CRFs, we measured the average response in each condition in a fixed window 3-9 s after stimulus onset (shaded gray area). (b) CRFs for the attended and unattended conditions for each ROI. The parameters that differed between conditions are shown in the lower right corner of each plot (* denotes *p <* 0.05, *** denotes *p <* 0.001; also see Table 1). Error bars shows ±1 bootstrapped SEM across subjects.

It is important to note that the measured CRFs did not clearly saturate, which can make it difficult to differentiate between a change in contrast gain and a change in response gain. However, we found that CRFs in each attention condition converged at high contrasts, particularly in V1 and V2. This pattern cannot be explained by a pure change in response gain, which gives us confidence that feature-based attention truly modulates contrast gain. Importantly, despite any uncertainty about whether feature-based attention modulates contrast gain or responses gain, our data provide clear evidence for a *multiplicative* effect of attention on BOLD CRFs.

### The effect of feature-based attention varies with voxel eccentricity preference

Recent work has found that CRF saturation varies with the eccentricity of voxel receptive fields, with CRFs saturating at lower contrasts (i.e. lower *C*_*50*_ values) for voxels with pRFs centered at lower eccentricities (i.e. closer to fovea) than for voxels that respond to the visual periphery (Vinke et al., 2022). Therefore, in an exploratory analysis, we examined whether the effect of feature-based attention varies with voxel eccentricity. We divided voxels in each visual area into three eccentricity bins (spanning 1.0–2.5°, 2.5–4.8°, and 4.8–8.5°) that roughly equated the number of voxels in each bin for each visual area. Figure 6 shows CRFs for each visual area as a function of voxel eccentricity preference (see Table 2 for Naka-Rushton parameter estimates). Like Vinke and colleagues (2022), we found that *C*_*50*_ values gradually increased with eccentricity. The contrast-gain attentional effect that we observed at the whole-ROI level was consistent across eccentricity bins (Table 2). Moreover, we also found that feature-based attention increased both contrast gain and response gain in V2 and V3 for the inner eccentricities, suggesting that the nature of attentional modulation varies with the eccentricity preference of neural populations. However, given the exploratory nature of this analysis, further work is necessary to test the robustness of this pattern.

**Table 2.** Naka-Rushton parameter estimates and resampling statistics results as a function of voxel eccentricity (corresponds to Figure 6).

| ROI | Eccentricity | Attended (mean ± bootstrapped SEM) | | | |
| --- | --- | --- | --- | --- | --- |
| | | Unattended (mean ± bootstrapped SEM) | | | |
|  |  | *p* value |  |  |  |
| | | $R_{\text{Max}}$ | $C_{50}$ | $b$ | $n$ |
| V1 | 1.0-2.5° | 0.76 ± 0.06 | 22.10 ± 2.81 | -0.34 ± 0.06 | 0.73 ± 0.10 |
| | | 0.68 ± 0.06 | 30.96 ± 4.18 | -0.33 ± 0.05 | 0.94 ± 0.11 |
| | | $p = 0.081$ | $p < 0.001^{***}$ | $p = 0.759$ | $p = 0.127$ |
| | 2.5-4.8° | 0.71 ± 0.06 | 28.19 ± 3.46 | -0.23 ± 0.05 | 0.87 ± 0.09 |
| | | 0.69 ± 0.06 | 42.05 ± 5.28 | -0.22 ± 0.04 | 1.27 ± 0.19 |
| | | $p = 0.435$ | $p < 0.001^{***}$ | $p = 0.913$ | $p < 0.001^{***}$ |
| | 4.8-8.5° | 0.68 ± 0.06 | 36.87 ± 3.07 | -0.16 ± 0.04 | 1.10 ± 0.09 |
| | | 0.67 ± 0.05 | 43.18 ± 3.29 | -0.17 ± 0.03 | 1.31 ± 0.15 |
| | | $p = 0.611$ | $p = 0.011^*$ | $p = 0.359$ | $p = 0.009^{**}$ |
| V2 | 1.0-2.5° | 0.41 ± 0.05 | 13.96 ± 2.46 | -0.23 ± 0.03 | 1.53 ± 0.43 |
| | | 0.33 ± 0.05 | 24.82 ± 6.02 | -0.23 ± 0.03 | 1.36 ± 0.50 |
| | | $p = 0.037^*$ | $p = 0.001^{**}$ | $p = 0.912$ | $p = 0.722$ |
| | 2.5-4.8° | 0.38 ± 0.04 | 25.24 ± 4.26 | -0.14 ± 0.03 | 1.23 ± 0.41 |
| | | 0.34 ± 0.04 | 41.07 ± 6.23 | -0.16 ± 0.03 | 1.48 ± 0.37 |
| | | $p = 0.146$ | $p < 0.001^{***}$ | $p = 0.278$ | $p = 0.419$ |
| | 4.8-8.5° | 0.37 ± 0.04 | 34.58 ± 4.27 | -0.09 ± 0.02 | 1.60 ± 0.27 |
| | | 0.37 ± 0.04 | 45.50 ± 3.31 | -0.11 ± 0.02 | 2.01 ± 0.35 |
| | | $p = 0.985$ | $p = 0.001^{**}$ | $p = 0.071$ | $p = 0.268$ |
| V3 | 1.0-2.5° | 0.35 ± 0.04 | 17.80 ± 2.56 | -0.14 ± 0.03 | 1.74 ± 0.42 |
| | | 0.27 ± 0.04 | 27.88 ± 4.84 | -0.14 ± 0.03 | 1.86 ± 0.51 |
| | | $p = 0.006^{**}$ | $p = 0.002^{**}$ | $p = 0.991$ | $p = 0.731$ |
| | 2.5-4.8° | 0.33 ± 0.03 | 25.49 ± 4.44 | -0.08 ± 0.02 | 1.47 ± 0.29 |
| | | 0.28 ± 0.03 | 39.48 ± 6.05 | -0.09 ± 0.02 | 1.73 ± 0.43 |
| | | $p = 0.019^*$ | $p = 0.001^{**}$ | $p = 0.559$ | $p = 0.495$ |
| | 4.8-8.5° | 0.35 ± 0.03 | 31.49 ± 4.48 | -0.08 ± 0.02 | 1.52 ± 0.23 |
| | | 0.34 ± 0.03 | 42.07 ± 4.92 | -0.10 ± 0.02 | 1.76 ± 0.32 |
| | | $p = 0.640$ | $p = 0.012^*$ | $p = 0.070$ | $p = 0.409$ |

\* denotes $p$ -values $< 0.05$ , \*\* denotes $p$ -values $< 0.01$ , and \*\*\* denotes $p$ -values $< 0.001$

**Figure 6.**
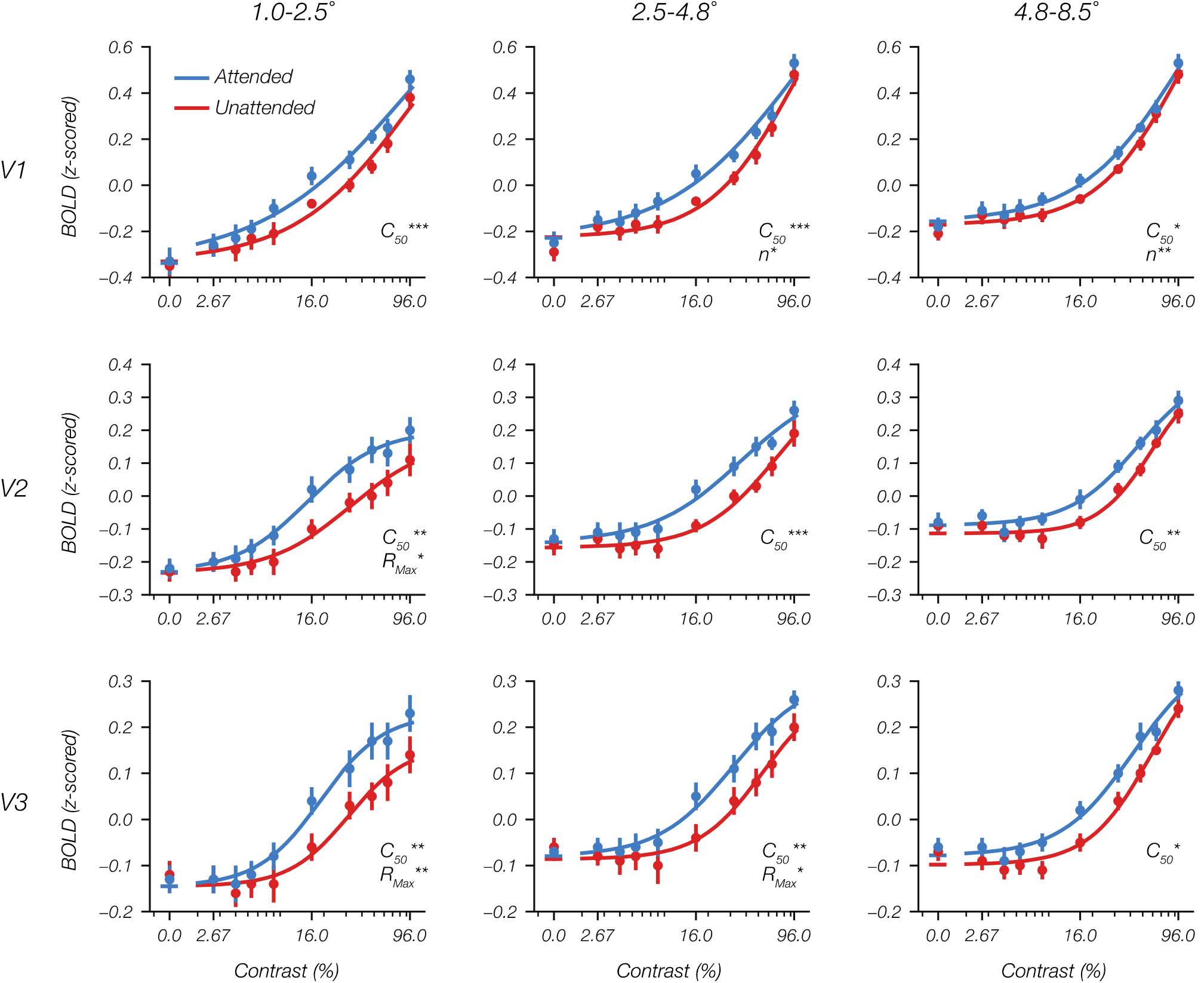
The effect of feature-based attention on probe-evoked CRFs as a function of voxel eccentricity preference. For each visual area (rows), we split voxels into bins based on eccentricity derived from pRF estimates (columns). The parameters that differed between conditions are shown in the lower right corner of each plot (* denotes *p <* 0.05, ** denotes *p <* 0.01, and *** denotes *p <* 0.001; also see Table 2). Error bars show ±1 bootstrapped SEM across subjects.

### Reconciling a contrast-gain effect with the normalization model of attention

Our finding that feature-based attention primarily increases contrast gain appears to be at odds with predictions from the normalization model of attention (NMA; Reynolds and Heeger, 2009), a prominent computational model of attention. Whereas we found that feature-based attention primarily increases contrast gain, Herrmann and colleagues (2012) showed that the NMA predicts that feature-based attention selectively increases response gain, leaving contrast gain unchanged, and reported psychophysical data that support this prediction. Thus, we turned to simulations using the NMA to investigate whether our contrast gain effect can be reconciled with the model (Fig. 7a). For both of the simulations reported below, we used Reynold and Heeger’s (2009) Matlab implementation of the NMA, which is available online (http://www.cns.nyu.edu/heegerlab/index.php?page=software&id=attentionModel). Table 3 summarizes the model parameters used in our simulations.

**Table 3.** NMA simulation model parameters

|  | Simulation 1 (Fig. 7b) | Simulation 2 (Fig. 7c) |
| --- | --- | --- |
| Orientation input(s) | **-45° + 45°** | **-45° only** |
| Attended orientation | -45° vs. 45° | -45° vs. 45° |
| Attentional field size |  |  |
| space | uniform | uniform |
| orientation | 30° | 30° |
| Stimulation field size |  |  |
| space | 5 (au) | 5 (au) |
| orientation | 20° | 20° |
| Suppressive field size |  |  |
| space | 20 (au) | 20 (au) |
| orientation | uniform | uniform |
| Measured neuron(s) | **tuned for -45°** | **Population average** |

The attentional field, stimulation field, and suppressive field were modeled as Gaussians, with size specified as their standard deviation (SD); rows where values differed between the two simulations are bolded; au = arbitrary units

**Figure 7.**
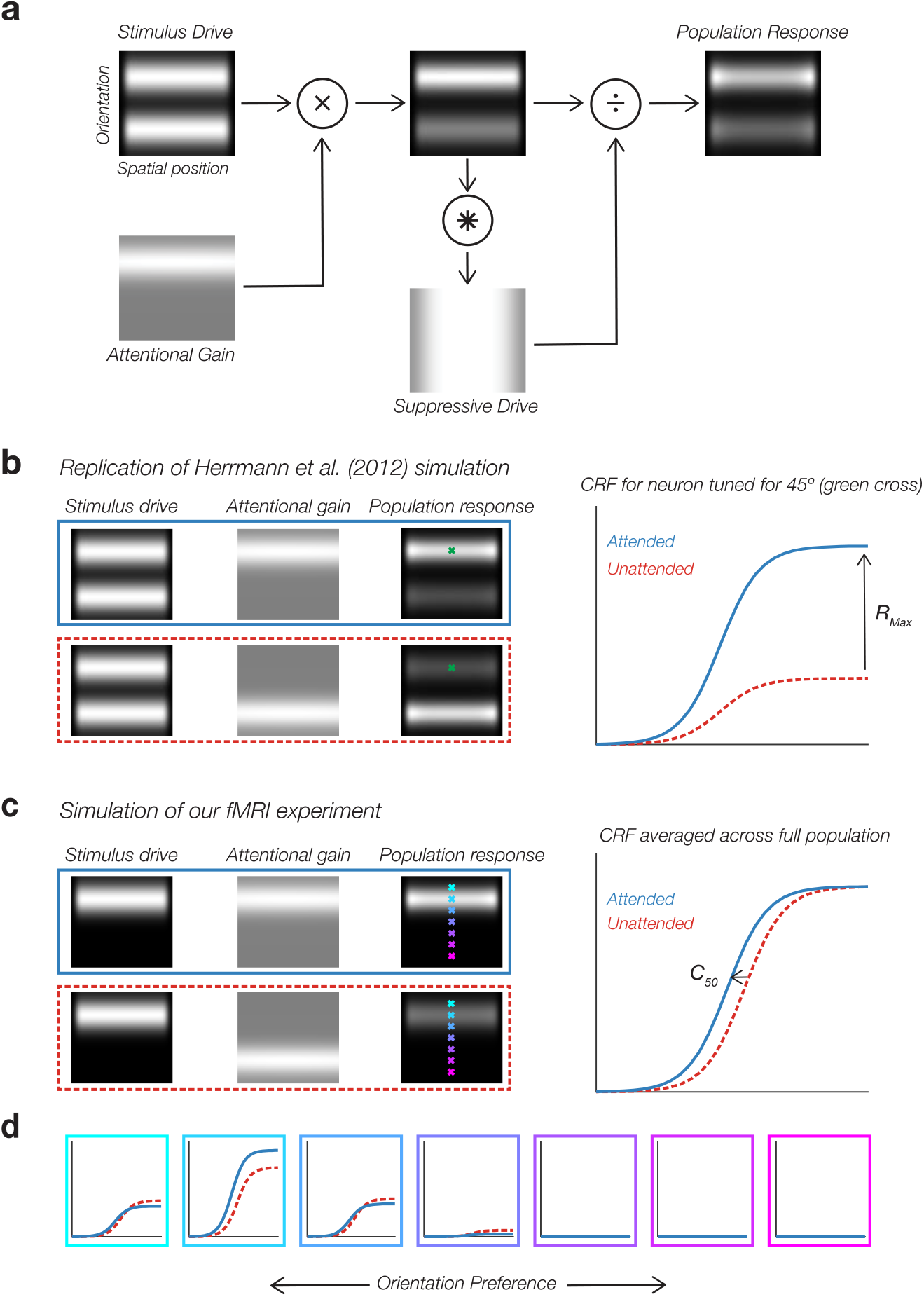
Normalization model of attention (NMA) and simulation results. (a) Schematic of the NMA (Reynolds & Heeger, 2009). The upper left panel shows the stimulus drive (brightness) as a function of the neuron’s receptive field center (horizontal axis) and orientation preference (vertical axis). In this example, two spatially overlapping stimuli of orthogonal orientations were presented to the model. The stimulus drive is multiplied by an attentional gain (lower left). The resulting product is convolved with a suppressive field to obtain suppressive drive. The final population response is calculated by dividing the product of stimulus drive and attentional gain by the suppressive drive. In our simulations, the suppressive field was Gaussian across space and uniform across orientation (see Table 3 for simulation parameters). (b) Replication of Herrmann et al. (2012) simulation. We presented two orthogonal orientations to the model (stimulus drive), and varied which orientation was attended (attentional gain), measuring the response of a neuron (green cross) when its preferred orientation was attended (blue line) or unattended (red dashed line). The predicted CRFs are shown on the right. Like Herrmann et al. (2012), we found that feature-based attention selectively increased response gain. (c) To model our own fMRI experiment, we presented a single probe orientation, and varied whether that orientation was attended or unattended. Because fMRI aggregates activity across neurons regardless of their orientation preference, we measured the average response across the entire orientation selective population. Under these conditions, we found that the NMA predicts that feature-based attention selectively increases contrast gain. (d) Probing further, we found that the effect of feature-based attention in our second simulation (b) varied as a function of the orientation preference in individual neurons. All neurons showed an increase in contrast gain. However, only neurons tuned for the probe orientation showed increased response gain. Neurons tuned for other orientations showed a *decrease* in response gain.

In our first simulation, we reproduced Herrmann et al.’s (2012) finding that feature-based attention selectively increases response gain (Fig. 7b). Following Herrmann et al. (2012), the stimulus input to the model was two orthogonal orientations, as was the case in their psychophysical experiment, and we measured the response for a simulated neuron that was tuned for one of the two presented orientations (marked by a green cross) when that orientation was attended or unattended (see Table 3 for all simulation parameters). Consistent with Herrmann et al.’s (2012) results, we found that feature-based attention selectively increased response gain under these conditions.

In our second simulation, we modeled the details of our own fMRI experiment (Fig. 7c). Our experiment differed from Herrmann and colleagues’ experiment in two important ways. First, whereas Herrmann et al. (2012) cued observers to attend one of two orthogonal orientations, in our study we presented a single orientation in the probe hemifield and manipulated the attended orientation in the other hemifield. To simulate our paradigm, we provided a single orientation as stimulus input, and we varied whether attentional gain was centered on that orientation or on the orthogonal orientation. Second, Herrmann et al. (2012) measured the response of neurons tuned for one of the two presented orientations when that orientation was attended or unattended. This decision is reasonable when modelling behavioral performance, which may largely depend on the gain of neurons tuned for the target feature. However, responses measured with fMRI aggregate across many neurons, regardless of their orientation selectivity. Thus, in our second simulation, we measured the average activity across the entire orientation-selective population. Apart from these two changes, the parameters of this simulation were otherwise identical to those used in the previous simulation (see Table 3). Under these conditions, we found that feature-based attention purely increased contrast gain (Fig. 7c).

It is striking that we found no evidence that feature-based attention increased response gain in our second simulation. However, the effect of attention predicted by the NMA can vary substantially across neurons with different tuning preferences (e.g. Hara et al., 2014). Thus, we examined the effect of feature-based attention across neurons with different orientation preferences (Fig. 7d). We found that feature-based attention uniformly increased contrast gain across neurons with different orientation preferences, but only increased response gain for neurons whose preferred orientation was close to the probe orientation. For neurons tuned for other orientations, feature-based attention *decreased* response gain. This decrease in response gain reflects the fact that these neurons receive little stimulus drive, but are subject to greater suppressive drive (which was uniform across orientation), when attention is directed to the probe orientation.

The results of these simulations show that our fMRI findings are broadly consistent with the NMA when the specific conditions of our experiment are modeled: pooled across the population, feature-based attention primarily increases contrast gain of CRFs. Thus, our fMRI results should not be taken as evidence against the NMA. That being said, we found some evidence for an increase in response gain for voxels responses preferring inner eccentricities (Fig. 6), and it remains unclear how this finding can be reconciled with the NMA. However, given the exploratory nature of our analysis examining how the effect of feature-based attention varied with eccentricity, we think it is important that this pattern is replicated before being considered too closely.

## Discussion

One primary way that attention improves neural coding is by increasing sensory gain, selectively amplifying stimulus-evoked neural responses (Hillyard et al., 1998; Carrasco, 2011). However, substantial evidence suggests that fMRI is not sensitive to these multiplicative effects of attention. Functional MRI studies that have examined the effect of spatial attention on CRFs have not observed the multiplicative effects that are seen in electrophysiological measures. Instead, attention produces an additive baseline shift in the BOLD signal (e.g. Buracas and Boynton, 2007; Murray, 2008; Sprague et al., 2018), calling into question what fMRI can tell us about how attention modulates sensory codes. In contrast with studies of spatial attention, a seminal study found that *feature-based attention* amplified the BOLD response to a probe stimulus when it matched the attended feature (Saenz et al., 2002), an apparent multiplicative attention effect that is difficult to explain if fMRI is not sensitive to modulation of sensory gain. However, no study to date has examined the effect of feature-based attention on the full CRF to definitively characterize the nature of this effect. In the current study, we provided that critical test, measuring the effect of feature-based attention on CRFs in early visual areas V1-V3. We found robust multiplicative effects of attention in early visual cortex. In all three areas, feature-based attention increased contrast gain, improving neural sensitivity. In V2 and V3 we also found some evidence for an increase in response gain, primarily in voxels that respond to inner eccentricities, increasing neural responsivity. Taken together, our results show that BOLD CRFs are multiplicatively modulated by feature-based attention.

Given the central role of fMRI in studying the neural mechanisms of attention, it is essential that we understand whether fMRI-BOLD is sensitive to multiplicative changes in sensory gain. Our results provide clear, positive evidence that fMRI is sensitive to these effects. However, our results stand in stark contrast with studies of spatial attention that have repeatedly found additive baseline shifts rather than multiplicative effects (e.g. Buracas and Boynton, 2007; Murray, 2008; Sprague et al., 2018; Itthipuripat et al., 2019). If the BOLD signal is sensitive to multiplicative changes in sensory gain, then why are these effects not seen in studies of spatial attention? Work by Hara and colleagues (2014) provides one possible answer to this discrepancy. Using the NMA, Hara and colleagues simulated the effect of spatial attention on population responses and found that the NMA predicts that the effect of spatial attention on the CRFs of simulated neurons depends on their tuning properties: spatial attention increased response gain for neurons that were well-tuned tuned for the feature value of the stimulus, but instead increased contrast gain for neurons tuned for other feature values. Hara and colleagues found that the net effect of these modulations on the population CRF was a near-additive effect. Thus, the additive effects of spatial attention reported in past fMRI studies may reflect the fact that the responses measured in these studies aggregated across neurons with different tuning preferences, rather than fMRI being inherently insensitive to multiplicative effects of attention. By contrast, in our simulations of feature-based attention, we found that the NMA predicts that feature-based attention, unlike spatial attention, will produce a multiplicative contrast-gain effect in the population response. Thus, feature-based attention may provide a clearer testbed than spatial attention for examining whether fMRI is sensitive to modulation of sensory gain.

Although Hara et al.’s (2014) simulations offer a plausible explanation for why fMRI studies of spatial attention have not detected multiplicative effects of attention, this account is yet to be empirically tested. Fortunately, recent advances in computational neuroimaging techniques provide an opportunity to test this account. Inverted encoding models (IEMs), for instance, allow researchers to reconstruct the activity of feature-selective channels (or neuronal populations) from voxelwise patterns of activity in visual cortex (Brouwer and Heeger, 2009, 2011). The IEM approach has proven to be a powerful tool for characterizing how attention modulates feature-selective population responses (e.g. Garcia et al., 2013; Sprague and Serences, 2013). Furthermore, this approach has been used to measure the effect of sensory manipulations (e.g. cross-orientation suppression) on the CRF of different orientation-selective channels (Brouwer and Heeger, 2011). Thus, the IEM approach provides an opportunity to test Hara and colleagues’ (2014) account that spatial attention increases response gain in the channel tuned for the stimulus value and increases contrast gain for channels tuned for other feature values.

Testing Hara et al.’s (2014) account is especially important considering a recent study by Itthipuripat and colleagues (2019), who examined the effect of spatial attention on CRFs measured with both fMRI and EEG in the same human observers performing the same attention task. Consistent with previous work, they found spatial attention produced a near-additive baseline shift in the fMRI-BOLD response, but increased response gain of EEG responses (e.g. the P1 component and steady-state-visual-evoked potentials), providing clear evidence that fMRI and EEG measurements are sensitive to different neural signals that are modulated by attention in different ways. Although Hara et al.’s (2014) account can explain the additive effect of attention measured with fMRI, Itthipuripat et al. (2019) pointed out that this account is difficult to reconcile with the EEG results because EEG, like fMRI, measures population-level activity, and would also be expected to show an additive shift. That being said, it is also possible that EEG may be particularly sensitive to neural signals that are primarily sensitive to changes in response gain, and much more work is needed to further understand why EEG and fMRI exhibit qualitatively distinct attentional effects.

## Conclusions

Attention improves sensory processing by increasing sensory gain, amplifying sensory responses evoked by attended stimuli. However, available evidence has suggested that the fMRI-BOLD signal is not sensitive to these multiplicative effects of attention. In the current study, we found that feature-based attention multiplicative scales CRFs in early visual cortex measured with fMRI, primarily increasing contrast gain on BOLD CRFs – an effect that is broadly consistent with predictions of how feature-based attention should modulate population neural responses derived from the NMA. Our results provide clear, positive evidence that fMRI is sensitive to multiplicative modulations of sensory gain. Given the central role of fMRI in the study of visual attention, an important direction for future work will be understanding the boundary conditions under which such multiplicative effects are observed.

## Acknowledgements

This work was supported by R01 EY28163 to SL, and a Boston University Center for Systems Neuroscience Postdoctoral Fellowship to JJF. This research was conducted at the Boston University Cognitive Neuroimaging Center, and involved the use of instrumentation supported by the NSF Major Research Instrumentation grant BCS-1625552. We acknowledge the University of Minnesota Center for Magnetic Resonance Research for use of the multiband-EPI pulse sequences. Data was analyzed on a high-performance computing cluster supported by ONR grant N00014-17-1-2304. We thank Shruthi Chakrapani for assistance with data collection, and members of the Ling Lab for helpful feedback on the manuscript.

## References

Albrecht DG, Hamilton DB (1982) Striate cortex of monkey and cat: contrast response function. Journal of Neurophysiology 48:217–237.

Andersson JLR, Skare S, Ashburner J (2003) How to correct susceptibility distortions in spin-echo echo-planar images: application to diffusion tensor imaging. NeuroImage 20:870–888.

Benson NC, Jamison KW, Arcaro MJ, Vu AT, Glasser MF, Coalson TS, Van Essen DC, Yacoub E, Ugurbil K, Winawer J, Kay K (2018) The Human Connectome Project 7 Tesla retinotopy dataset: Description and population receptive field analysis. Journal of Vision 18:23.

Boynton GM, Demb JB, Glover GH, Heeger DJ (1999) Neuronal basis of contrast discrimination. Vision Research 39:257–269.

Brainard DH (1997) The Psychophysics Toolbox. Spatial Vision 10:433–436.

Brouwer GJ, Heeger DJ (2009) Decoding and reconstructing color from responses in human visual cortex. Journal of Neuroscience 29:13992–14003.

Brouwer GJ, Heeger DJ (2011) Cross-orientation suppression in human visual cortex. Journal of neurophysiology 106:2108–2119.

Buracas GT, Boynton GM (2007) The effect of spatial attention on contrast response functions in human visual cortex. Journal of Neuroscience 27:93–97.

Carrasco M (2011) Visual attention: The past 25 years. Vision Research 51:1484–1525.

Cauley SF, Polimeni JR, Bhat H, Wald LL, Setsompop K (2014) Interslice leakage artifact reduction technique for simultaneous multislice acquisitions. Magnetic Resonance in Medicine 72:93–102.

Chelazzi L, Miller EK, Duncan J, Desimone R (1993) A neural basis for visual search in inferior temporal cortex. Nature 363:345–347.

Dale AM (1999) Optimal experimental design for event-related fMRI. Human Brain Mapping 8:109–114.

Duncan J, Ward R, Shapiro K (1994) Direct measurement of attentional dwell time in human vision. Nature 369:313–315.

Feinberg DA, Moeller S, Smith SM, Auerbach E, Ramanna S, Glasser MF, Miller KL, Ugurbil K, Yacoub E (2010) Multiplexed echo planar imaging for sub-second whole brain fMRI and fast diffusion imaging. PLOS ONE 5:e15710.

Fischl B (2012) FreeSurfer. NeuroImage 62:774–781.

Foster JJ, Thyer W, Wennberg JW, Awh E (2021) Covert attention increases the gain of stimulus-evoked population codes. J Neurosci 41:1802–1815.

Garcia JO, Srinivasan R, Serences JT (2013) Near-real-time feature-selective modulations in human cortex. Current Biology 23:515–522.

Gardner JL, Sun P, Waggoner RA, Ueno K, Tanaka K, Cheng K (2005) Contrast adaptation and representation in human early visual cortex. Neuron 47:607–620.

Greve DN, Fischl B (2009) Accurate and robust brain image alignment using boundary-based registration. NeuroImage 48:63–72.

Griswold MA, Jakob PM, Heidemann RM, Nittka M, Jellus V, Wang J, Kiefer B, Haase A (2002) Generalized autocalibrating partially parallel acquisitions (GRAPPA). Magnetic Resonance in Medicine 47:1202–1210.

Hara Y, Pestilli F, Gardner JL (2014) Differing effects of attention in single-units and populations are well predicted by heterogeneous tuning and the normalization model of attention. Front Comput Neurosci 8.

Herrmann K, Heeger DJ, Carrasco M (2012) Feature-based attention enhances performance by increasing response gain. Vision Research 74:10–20.

Hesterberg T (2011) Bootstrap. WIREs Computational Statistics 3:497–526.

Hillyard SA, Vogel EK, Luck SJ (1998) Sensory gain control (amplification) as a mechanism of selective attention: electrophysiological and neuroimaging evidence. Philosophical transactions of the Royal Society of London Series B, Biological Sciences 353:1257–1270.

Itthipuripat S, Ester EF, Deering S, Serences JT (2014a) Sensory gain outperforms efficient readout mechanisms in predicting attention-related improvements in behavior. J Neurosci 34:13384–13398.

Itthipuripat S, Garcia JO, Rungratsameetaweemana N, Sprague TC, Serences JT (2014b) Changing the spatial scope of attention alters patterns of neural gain in human cortex. Journal of Neuroscience 34:112–123.

Itthipuripat S, Sprague TC, Serences JT (2019) Functional MRI and EEG index complementary attentional modulations. Journal of Neuroscience 39:6162–6179.

Kay KN, Winawer J, Mezer A, Wandell BA (2013) Compressive spatial summation in human visual cortex. Journal of Neurophysiology 110:481–494.

Kim YJ, Grabowecky M, Paller KA, Muthu K, Suzuki S (2007) Attention induces synchronization-based response gain in steady-state visual evoked potentials. Nature Neuroscience 10:117–125.

Li X, Lu Z-L, Tjan BS, Dosher BA, Chu W (2008) Blood oxygenation level-dependent contrast response functions identify mechanisms of covert attention in early visual areas. Proceedings of the National Academy of Sciences 105:6202–6207.

Martínez-Trujillo JC, Treue S (2002) Attentional modulation strength in cortical area MT depends on stimulus contrast. Neuron 35:365–370.

McAdams CJ, Maunsell JH (1999) Effects of attention on orientation-tuning functions of single neurons in macaque cortical area V4. Journal of Neuroscience 19:431–441.

Moeller S, Yacoub E, Olman CA, Auerbach E, Strupp J, Harel N, Uğurbil K (2010) Multiband multislice GE-EPI at 7 tesla, with 16-fold acceleration using partial parallel imaging with application to high spatial and temporal whole-brain fMRI. Magnetic Resonance in Medicine 63:1144–1153.

Morgan ST, Hansen JC, Hillyard SA (1996) Selective attention to stimulus location modulates the steady-state visual evoked potential. Proceedings of the National Academy of Sciences 93:4770–4774.

Murray SO (2008) The effects of spatial attention in early human visual cortex are stimulus independent. Journal of vision 8:2.1-11.

Ohzawa I, Sclar G, Freeman RD (1982) Contrast gain control in the cat visual cortex. Nature 298:266–268.

Pelli DG (1997) The VideoToolbox software for psychophysics: transforming numbers into movies. Spatial Vision 10:437–442.

Pestilli F, Carrasco M, Heeger DJ, Gardner JL (2011) Attentional enhancement via selection and pooling of early sensory responses in human visual cortex. Neuron 72:832–846.

Raymond JE, Shapiro KL, Arnell KM (1992) Temporary suppression of visual processing in an RSVP task: An attentional blink? Journal of Experimental Psychology: Human Perception and Performance 18:849–860.

Reuter M, Rosas HD, Fischl B (2010) Highly accurate inverse consistent registration: A robust approach. NeuroImage 53:1181–1196.

Reynolds JH, Heeger DJ (2009) The normalization model of attention. Neuron 61:168–185.

Reynolds JH, Pasternak T, Desimone R (2000) Attention increases sensitivity of V4 neurons. Neuron 26:703–714.

Saenz M, Buracas GT, Boynton GM (2002) Global effects of feature-based attention in human visual cortex. Nature Neuroscience 5:631–632.

Sclar G, Lennie P, DePriest DD (1989) Contrast adaptation in striate cortex of macaque. Vision Research 29:747–755.

Serences JT, Boynton GM (2007) Feature-based attentional modulations in the absence of direct visual stimulation. Neuron 55:301–312.

Setsompop K, Gagoski BA, Polimeni JR, Witzel T, Wedeen VJ, Wald LL (2012) Blipped-controlled aliasing in parallel imaging for simultaneous multislice echo planar imaging with reduced g-factor penalty. Magnetic Resonance in Medicine 67:1210–1224.

Smith SM, Jenkinson M, Woolrich MW, Beckmann CF, Behrens TEJ, Johansen-Berg H, Bannister PR, De Luca M, Drobnjak I, Flitney DE, Niazy RK, Saunders J, Vickers J, Zhang Y, De Stefano N, Brady JM, Matthews PM (2004) Advances in functional and structural MR image analysis and implementation as FSL. NeuroImage 23:S208–S219.

Somers DC, McMains SA (2005) Spatially-specific attentional modulation revealed by fMRI. In: Neurobiology of Attention, pp 377–382. Academic Press.

Sprague TC, Itthipuripat S, Vo V, Serences JT (2018) Dissociable signatures of visual salience and behavioral relevance across attentional priority maps in human cortex. Journal of Neurophysiology:1–26.

Sprague TC, Serences JT (2013) Attention modulates spatial priority maps in the human occipital, parietal and frontal cortices. Nature Neuroscience 16:1879–1887.

Stokes M, Thompson R, Nobre AC, Duncan J (2009) Shape-specific preparatory activity mediates attention to targets in human visual cortex. Proceedings of the National Academy of Sciences of the United States of America 106:19569–19574.

Tootell RB, Hadjikhani N, Hall EK, Marrett S, Vanduffel W, Vaughan JT, Dale AM (1998) The retinotopy of visual spatial attention. Neuron 21:1409–1422.

Treue S, Martinez-Trujillo JC (1999) Feature-based attention influences motion processing gain in macaque visual cortex. Nature 399:575–579.

van der Kouwe AJW, Benner T, Salat DH, Fischl B (2008) Brain morphometry with multiecho MPRAGE. NeuroImage 40:559–569.

van Voorhis S, Hillyard SA (1977) Visual evoked potentials and selective attention to points in space. Perception & Psychophysics 22:54–62.

Vinke LN, Bloem IM, Ling S (2022) Saturating nonlinearities of contrast response in human visual cortex. J Neurosci 42:1292–1302.

Virtanen P et al. (2020) SciPy 1.0: fundamental algorithms for scientific computing in Python. Nat Methods 17:261–272.

Williford T, Maunsell JHR (2006) Effects of spatial attention on contrast response functions in Macaque area V4. Journal of Neurophysiology 96:40–54.

Xu J, Moeller S, Auerbach EJ, Strupp J, Smith SM, Feinberg DA, Yacoub E, Uğurbil K (2013) Evaluation of slice accelerations using multiband echo planar imaging at 3T. NeuroImage 83:991–1001.

Yeo BTT, Krienen FM, Sepulcre J, Sabuncu MR, Lashkari D, Hollinshead M, Roffman JL, Smoller JW, Zöllei L, Polimeni JR, Fischl B, Liu H, Buckner RL (2011) The organization of the human cerebral cortex estimated by intrinsic functional connectivity. Journal of Neurophysiology 106:1125–1165.

